# scSampler: fast diversity-preserving subsampling of large-scale single-cell transcriptomic data

**DOI:** 10.1101/2022.01.15.476407

**Authors:** Dongyuan Song, Nan Miles Xi, Jingyi Jessica Li, Lin Wang

## Abstract

**Summary:** The number of cells measured in single-cell transcriptomic data has grown fast in recent years. For such large-scale data, subsampling is a powerful and often necessary tool for exploratory data analysis. However, the easiest random subsampling is not ideal from the perspective of preserving rare cell types. Therefore, diversity-preserving subsampling is required for fast exploration of cell types in a large-scale dataset. Here we propose scSampler, an algorithm for fast diversity-preserving subsampling of single-cell transcriptomic data. Using simulated and real data, we show that scSampler consistently outperforms existing subsampling methods in terms of both the computational time and the Hausdorff distance between the full and subsampled datasets.

**Availability:** scSampler is implemented in Python and is published under the MIT source license. It can be installed by `pip install scsampler` and used with the Scanpy pipline. The code is available on GitHub: https://github.com/SONGDONGYUAN1994/scsampler.

## 1 Introduction

Single-cell RNA sequencing (scRNA-seq) technologies have undergone rapid development in recent years. A remarkable achievement is the generation of large-scale datasets, and there are even datasets containing a million cells (See Supplementary Materials Table S1). Such massive scRNA-seq datasets have impeded exploratory data analysis (e.g., visualization) on standard computers.

An intuitive solution to this “big data” challenge is to subsample (downsample) a large-scale dataset, i.e., to select a subset of representative cells. Random subsampling is fast and unbiased, and it has been implemented in popular pipelines such as Seurat [1] and Scanpy [2]. However, random subsampling may miss rare cell types and is thus not ideal for preserving the transcriptome diversity. To overcome this drawback of random subsampling, Hie et al. proposed the first algorithm Geosketch for “intelligently selecting a subset of single cells”, which they called “sketching” [3]. Geosketch aims to evenly sample cells across the transcriptome space by minimizing the Hausdorff distance between the subsample and the original sample (i.e., the large-scale dataset) (**Supplementary Materials** S3.1). In the follow-up algorithm Hopper, the authors improved the performance of Geosketch in terms of minimizing the Hausdorff distance. Moreover, prior to Geosketch and Hopper and outside of the single-cell field, this “intelligent subsampling” problem has been well studied in the field of computer experiment design, in which the “space-filling designs” implement the idea of even subsampling across the transcriptomic space [4]. The most popular space-filling designs are the minimax distance design and the maximin distance design [5]. Geosketch and Hopper conceptually belong to the minimax distance design, which, however, is much more computationally intensive than the maximin distance design [5]. Here we propose scSampler, a Python package for fast diversity-preserving subsampling of large-scale single-cell transcriptomic data. By “diversity-preserving sampling,” scSampler implements the maximin distance design to make cells in the subsample as separative as possible. Using 8 simulated datasets and 10 real datasets, we show that scSampler outperforms existing subsampling methods in minimizing the Hausdorff distance between the subsample and the original sample. Moreover, scSampler is fast and scalable for million-level data.

## 2 Implementation

scSampler is implemented in Python and can be installed by `pip install scsampler`. The input is a matrix or an anndata object from scanpy pipeline. Denote the input matrix by **X** ∈ ℝ^*n×p*^, whose columns correspond to *p* features (by default, top *p* PCs from a cell-by-gene log(count + 1) matrix and scaled to [0, 1]) and whose rows correspond to *n* cells. Therefore, **X** can also represent a set *Χ* = {*x*_1_, …, *x*_*n*_}, where *x*_*i*_ ∈ ℝ^*p*^. Our goal is to find a size *n*_*s*_ subset *Χ*_*s*_ ⊂ *Χ*, which satisfies:

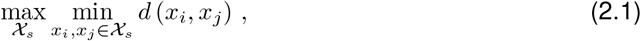

where *d*(·, ·) is the Euclidean distance. The optimality in (2.1) can be achieved by minimizing a scalar loss function:

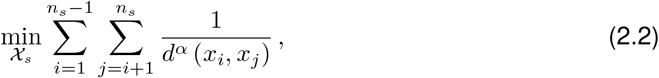

for a sufficiently large *α* [6]. It is found by [7] and in our numerical results that *α* = 4*p* is big enough and keeps the algorithm numerically stable, so we set *α* = 4*p* by default. For computational efficiency, scSampler can randomly split the original sample into subsets and perform subsampling on each subset. The detailed optimization algorithm is in **Supplementary Materials** S1.

## 3 Results

To comprehensively benchmark scSampler—three variants: scSampler-sp1 (no sample splitting; the slowest), scSampler-sp4 (splitting the sample into 4 subsets), and scSampler-sp16 (splitting the sample into 16 subsets; the fastest)—against random sampling and two state-of-the-art subsampling methods, Geosketch and Hopper, we use the scRNA-seq simulator Splatter [8] to generate 8 simulated datasets and collect 10 real datasets (See **Supplementary Materials** Table S1). On each dataset, we subsample 1000, 3000, 5000 and 10000 cells using each subsampling method. Fig. 1a shows an example that illustrates the difference between random subsampling and scSampler: compared to random sampling, scSampler selects more cells from small cell clusters. Quantitatively, we compare subsampling methods by two measures: (1) the Hausdorff distance between the subsample and the original sample, (2) computation time, both of which are better if smaller (Fig. 1b, **Supplementary Materials** Table S3-S4). Fig. 1b summarizes the performance of subsampling methods in the two measures. Notably, scSampler-sp1 consistently yields the smallest Hausdorff distances across all datasets and all subsample sizes. Moreover, scSampler is fast: on the largest cortex dataset (more than 1 million cells), scSampler-sp1 finishes in 15 minutes, and scSampler-sp16 takes only 1 minute and still outperforms Geosketch and Hopper by achieving a lower Hausdorff distance. Fig. 1c shows that scSampler is consistently ranked the top (smaller ranks are better) across the 18 datasets.

**Figure 1:**
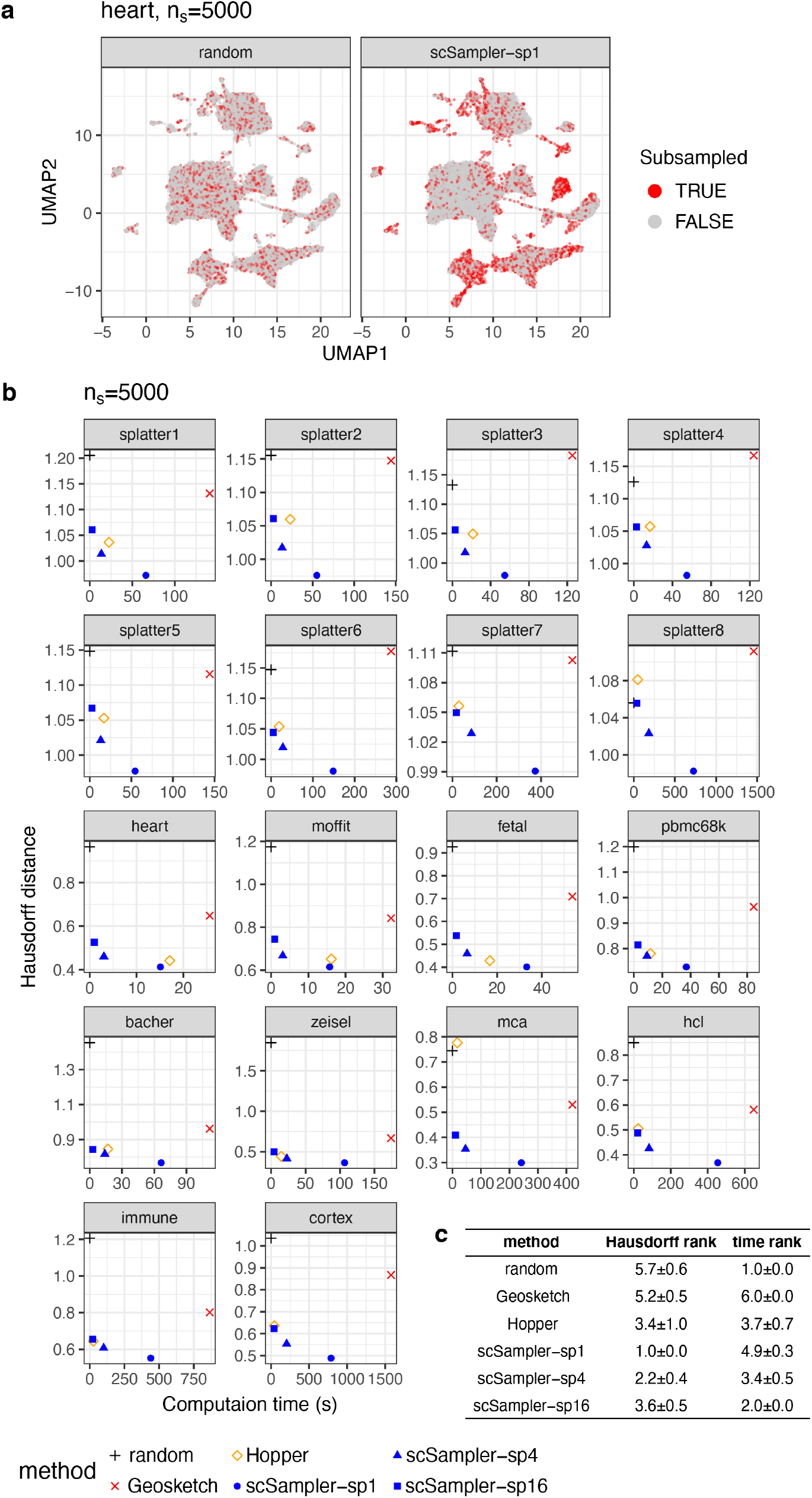
Benchmarking scSampler against other subsampling methods. (a) UMAP visualization of selected cells in the original data by random subsampling and scSampler-sp1, respectively. (b) Scatter plots of Hausdorff distance against computation time. (c) Summary of the performance of each methods. The table shows the mean ranks and standard deviations across all datasets and subsample sizes.

To verify if rare cell types are better captured by scSampler than other methods, we calculate the Gini coefficient of cell type proportions in each subsample; a smaller Gini coefficient indicates more balanced cell types (**Supplementary Materials** S3.3). In more than 60% of the combinations of 18 datasets and 4 subsmaple sizes, the fastest scSampler-sp16 leads to the smallest Gini coefficient (**Supplementary Materials** Table S5). Considering that the real datasets may not have accurately annotated cell types, we examine the simulated datasets and find that scSampler-sp16 leads to the smallest Gini coefficient in 90% of the combinations of 8 simulated datasets and 4 sample sizes, confirming that scSampler well preserves rare cell types.

## References

[1] Rahul Satija, Jeffrey A Farrell, David Gennert, Alexander F Schier, and Aviv Regev. Spatial reconstruction of single-cell gene expression data. Nature biotechnology, 33(5):495–502, 2015.

[2] F Alexander Wolf, Philipp Angerer, and Fabian J Theis. Scanpy: large-scale single-cell gene expression data analysis. Genome biology, 19(1):1–5, 2018.

[3] Brian Hie, Hyunghoon Cho, Benjamin DeMeo, Bryan Bryson, and Bonnie Berger. Geometric sketching compactly summarizes the single-cell transcriptomic landscape. Cell systems, 8(6):483–493, 2019.

[4] V Roshan Joseph. Space-filling designs for computer experiments: A review. Quality Engineering, 28(1):28–35, 2016.

[5] Mark E Johnson, Leslie M Moore, and Donald Ylvisaker. Minimax and maximin distance designs. Journal of statistical planning and inference, 26(2):131–148, 1990.

[6] Max D Morris and Toby J Mitchell. Exploratory designs for computational experiments. Journal of statistical planning and inference, 43(3):381–402, 1995.

[7] V Roshan Joseph, Tirthankar Dasgupta, Rui Tuo, and CF Jeff Wu. Sequential exploration of complex surfaces using minimum energy designs. Technometrics, 57(1):64–74, 2015.

[8] Luke Zappia, Belinda Phipson, and Alicia Oshlack. Splatter: simulation of single-cell rna sequencing data. Genome biology, 18(1):1–15, 2017.

