## Supplementary Materials for "scSampler: fast diversity-preserving subsampling of large-scale single-cell transcriptomic data"

### S1 Algorithm

**Algorithm 1** scSampler: sequential selection

**Input:** a sample (dataset) $\mathcal{X}$, a required subsample size $n_s$, and a split fold $B$
**Output:** a subsample $\mathcal{X}_s$
Evenly split $\mathcal{X}$ into $B$ random subsets $\mathcal{X}^1, \cdots, \mathcal{X}^b \cdots, \mathcal{X}^B$
**for** each $\mathcal{X}^b$ **do**
  $m = 1$
  Randomly select a point $x_1$ from $\mathcal{X}^b$
  Initialize $\mathcal{X}_s^b$ as a subsample with a single point $x_1$ and remove $x_1$ from $\mathcal{X}^b$
  **for** each $x \in \mathcal{X}^b$ **do**
    Compute $\Delta(x|\mathcal{X}_s^b) = 1/d^\alpha(x_1, x)$
  **end for**
  **while** $m < \lceil n_s/B \rceil$ **do**
    Find $x_{m+1}$ by (S1)
    Include $x_{m+1}$ in $\mathcal{X}_s^b$ and remove it from $\mathcal{X}^b$
    **for** each $x \in \mathcal{X}^b$ **do**
      Update $\Delta(x|\mathcal{X}_s^b) \leftarrow \Delta(x|\mathcal{X}_s^b) + 1/d^\alpha(x_{m+1}, x)$
    **end for**
    $m \leftarrow m + 1$
  **end while**
**end for**
$\mathcal{X}_s = \cup_{b=1}^B \mathcal{X}_s^b$

#### S1.1 The sequential criterion

We want to generate a subsample, denoted by $\mathcal{X}_s^b$, with $n_0 = \lceil n_s/B \rceil$ points from each split sample $\mathcal{X}^b$, $b = 1, \ldots, B$. We start with a random point $x_1$ and select $x_2, \ldots, x_{n_0}$ sequentially. Suppose we have already selected $m$ points. Then the $(m+1)$th point is obtained as

$$\begin{aligned} x_{m+1} &= \arg\min_{x \in \mathcal{X}^b} \left\{ \sum_{i=1}^{m-1} \sum_{j=i+1}^{m} \frac{1}{d^\alpha\left(x_i, x_j\right)} + \sum_{i=1}^{m} \frac{1}{d^\alpha\left(x_i, x\right)} \right\} \\ &= \arg\min_{x \in \mathcal{X}^b} \Delta(x|\mathcal{X}_s^b), \end{aligned} \quad \text{(S1)}$$

where $\alpha = 4p$ and

$$\Delta(x|\mathcal{X}_s^b) = \sum_{i=1}^{m} \frac{1}{d^\alpha\left(x_i, x\right)}.$$

#### S2 Datasets used in the paper

The datasets used in the paper were all stored in the `anndata` format of `Scanpy` pipeline. We followed the `Scanpy` tutorial to preprocess all datasets, including gene filtering (expressed in more than 3 cells), calculating mitochondrion mRNA, normalizing total counts into 10000, log(count + 1) transformation, selecting highly variable genes (HVGs), regressing out mitochondrion mRNA and cell total counts, scaling data and PCA (top 50 PCs). The input data for all subsampling methods were the top 50 PCs, each of which was scaled to $[0, 1]$ for a fair comparison of the Hausdorff distance. The Hausdorff distance in the nonscaled PC space can be computed by simply scaling back the selected subsample to the nonscaled PC space.

The simulated datasets were generated by R package `Splatter` ( [Zappia et al., 2017]). The basic parameters were group probability 0.01, 0.09, 0.2, 0.3, or 0.4 and DE probability 0.3, 0.1, 0.2, 0.01, or 0.1. For datasets splatter1–splatter8, we multiplied the DE probability by a real number $\lambda$ to control the distances between cell types.

Table S1: Overview of datasets used in the study

| dataset | # of cells | # of HVGs | # of cell types | $\lambda$ (splatter1–splatter8) or reference |
|---|---|---|---|---|
| splatter1 | 100000 | 529 | 5 | 0.1 |
| splatter2 | 100000 | 524 | 5 | 0.1 |
| splatter3 | 100000 | 515 | 5 | 0.15 |
| splatter4 | 100000 | 497 | 5 | 0.2 |
| splatter5 | 100000 | 500 | 5 | 0.25 |
| splatter6 | 200000 | 524 | 5 | 0.2 |
| splatter7 | 500000 | 510 | 5 | 0.2 |
| splatter8 | 1000000 | 517 | 5 | 0.2 |
| heart | 29552 | 2350 | 32 | [McLellan et al., 2020] |
| moffit | 30332 | 1729 | 12 | [Moffitt et al., 2018] |
| fetal | 61006 | 3330 | 32 | [Vento-Tormo et al., 2018] |
| pbmc68k | 68579 | 1488 | 10 | [Zheng et al., 2017] |
| bacher | 104417 | 803 | 6 | [Bacher et al., 2020] |
| zeisel | 160796 | 3389 | 39 | [Zeisel et al., 2015] |
| mca | 333778 | 3587 | 52 | [Han et al., 2018] |
| hcl | 599926 | 1864 | 63 | [Han et al., 2020] |
| immune | 606606 | 1566 | 34 | [Papatheodorou et al., 2020] |
| cortex | 1089022 | 2124 | 43 | [Yao et al., 2021] |

### S3 Metrics used in the study

#### S3.1 Hausdorff distance

The Hausdorff distance $d_{\mathrm{H}}$ measures the distance from the subsample to the original data. It is defined as:

$$d_{\mathrm{H}}(X_s, X) = \max_{x \in \mathcal{X}} \min_{x_s \in \mathcal{X}_s} d(x, x_s)\,, \tag{S2}$$

where $d$ is the Euclidean distance by default. Intuitively, a small Hausdorff distance means that all data points in the original sample can be represented well in the subsample. The Hausdorff distance is used as the primary metric for evaluating the quality of the subsamples.

#### S3.2 Computation time

All methods were running on a server with Ubuntu 20.04 system and used 1 core with 2.5G HZ. The times were recorded by Python `time` module.

#### S3.3 Gini coefficient

The Gini coefficient measures the inequality of cell types. A smaller Gini coefficient means that the cell types are more balanced in a sample. Here we denote the subsample as $X_s$, and the cell type proportions in $X_s$ are $p_1, \cdots, p_c, \cdots, p_C$. Note that the $C$ is the total number of cell types in the original data. Therefore, if a cell $c$ type is missed in the subsample, its proportion $p_c = 0$. Then the Gini coefficient $G$ is calculated as:

$$G = \frac{\sum_{c=1}^{C}(2c - C - 1)p_{(c)}}{C\sum_{c=1}^{C} p_{(c)}}\,, \tag{S3}$$

where $p_{(c)}$ is the $c$-th smallest cell type proportion. The code for calculation is from `https://github.com/oliviaguest/gini`.

#### S4 Performance of all subsampling methods

Table S3: The Hausdorff distances between subsamples and original samples

| dataset | random | Geosketch | Hopper | scSampler-sp1 | scSampler-sp4 | scSampler-sp16 | subsample size |
|---|---|---|---|---|---|---|---|
| splatter1 | 1.193 | 1.194 | 1.165 | 1.106 | 1.155 | 1.190 | 1000 |
| splatter2 | 1.239 | 1.225 | 1.182 | 1.112 | 1.165 | 1.182 | 1000 |
| splatter3 | 1.187 | 1.245 | 1.166 | 1.125 | 1.156 | 1.188 | 1000 |
| splatter4 | 1.226 | 1.230 | 1.180 | 1.125 | 1.187 | 1.227 | 1000 |
| splatter5 | 1.220 | 1.183 | 1.168 | 1.116 | 1.167 | 1.188 | 1000 |
| splatter6 | 1.158 | 1.182 | 1.173 | 1.111 | 1.158 | 1.154 | 1000 |
| splatter7 | 1.142 | 1.188 | 1.146 | 1.110 | 1.151 | 1.185 | 1000 |
| splatter8 | 1.123 | 1.161 | 1.141 | 1.096 | 1.130 | 1.145 | 1000 |
| heart | 1.032 | 0.864 | 0.648 | 0.579 | 0.660 | 0.752 | 1000 |
| moffit | 1.329 | 1.006 | 0.873 | 0.792 | 0.883 | 0.951 | 1000 |
| fetal | 1.032 | 0.808 | 0.631 | 0.558 | 0.650 | 0.778 | 1000 |
| pbmc68k | 1.299 | 1.206 | 1.001 | 0.922 | 0.984 | 1.024 | 1000 |
| bacher | 1.479 | 1.137 | 1.018 | 0.933 | 1.000 | 1.051 | 1000 |
| zeisel | 2.058 | 0.880 | 0.749 | 0.499 | 0.610 | 0.844 | 1000 |
| mca | 1.004 | 0.894 | 0.949 | 0.430 | 0.544 | 0.687 | 1000 |
| hcl | 0.922 | 0.849 | 0.827 | 0.502 | 0.596 | 0.719 | 1000 |
| immune | 1.963 | 1.001 | 0.863 | 0.696 | 0.779 | 0.903 | 1000 |
| cortex | 1.910 | 1.026 | 0.925 | 0.627 | 0.741 | 0.881 | 1000 |
| splatter1 | 1.208 | 1.168 | 1.077 | 1.016 | 1.061 | 1.093 | 3000 |
| splatter2 | 1.156 | 1.165 | 1.100 | 1.021 | 1.067 | 1.097 | 3000 |
| splatter3 | 1.152 | 1.162 | 1.094 | 1.024 | 1.062 | 1.099 | 3000 |
| splatter4 | 1.153 | 1.167 | 1.106 | 1.029 | 1.075 | 1.108 | 3000 |
| splatter5 | 1.171 | 1.152 | 1.091 | 1.022 | 1.073 | 1.116 | 3000 |
| splatter6 | 1.147 | 1.138 | 1.085 | 1.022 | 1.062 | 1.102 | 3000 |
| splatter7 | 1.112 | 1.129 | 1.097 | 1.028 | 1.076 | 1.088 | 3000 |
| splatter8 | 1.078 | 1.157 | 1.071 | 1.021 | 1.060 | 1.092 | 3000 |
| heart | 0.964 | 0.718 | 0.498 | 0.462 | 0.520 | 0.597 | 3000 |
| moffit | 1.174 | 0.904 | 0.716 | 0.670 | 0.727 | 0.789 | 3000 |
| fetal | 0.968 | 0.661 | 0.487 | 0.446 | 0.509 | 0.567 | 3000 |
| pbmc68k | 1.226 | 1.058 | 0.855 | 0.791 | 0.842 | 0.873 | 3000 |
| bacher | 1.454 | 1.007 | 0.912 | 0.817 | 0.870 | 0.923 | 3000 |
| zeisel | 1.844 | 0.682 | 0.486 | 0.400 | 0.465 | 0.549 | 3000 |
| mca | 0.785 | 0.561 | 0.683 | 0.334 | 0.408 | 0.486 | 3000 |
| hcl | 0.850 | 0.643 | 0.595 | 0.404 | 0.471 | 0.556 | 3000 |
| immune | 1.226 | 0.842 | 0.710 | 0.595 | 0.657 | 0.708 | 3000 |
| cortex | 1.048 | 0.968 | 0.706 | 0.527 | 0.605 | 0.689 | 3000 |
| splatter1 | 1.205 | 1.131 | 1.036 | 0.972 | 1.014 | 1.061 | 5000 |
| splatter2 | 1.155 | 1.147 | 1.060 | 0.976 | 1.017 | 1.061 | 5000 |
| splatter3 | 1.133 | 1.183 | 1.049 | 0.979 | 1.017 | 1.056 | 5000 |
| splatter4 | 1.126 | 1.167 | 1.057 | 0.982 | 1.028 | 1.057 | 5000 |
| splatter5 | 1.148 | 1.116 | 1.053 | 0.977 | 1.021 | 1.067 | 5000 |
| splatter6 | 1.147 | 1.178 | 1.054 | 0.980 | 1.019 | 1.044 | 5000 |
| splatter7 | 1.112 | 1.103 | 1.056 | 0.991 | 1.029 | 1.050 | 5000 |
| splatter8 | 1.056 | 1.112 | 1.081 | 0.982 | 1.023 | 1.056 | 5000 |
| heart | 0.964 | 0.648 | 0.442 | 0.413 | 0.460 | 0.526 | 5000 |
| moffit | 1.174 | 0.842 | 0.652 | 0.615 | 0.667 | 0.745 | 5000 |

| | | | | | | | |
|---|---|---|---|---|---|---|---|
| fetal | 0.926 | 0.709 | 0.429 | 0.401 | 0.459 | 0.538 | 5000 |
| pbmc68k | 1.199 | 0.964 | 0.781 | 0.729 | 0.771 | 0.815 | 5000 |
| bacher | 1.454 | 0.962 | 0.846 | 0.767 | 0.817 | 0.843 | 5000 |
| zeisel | 1.844 | 0.668 | 0.443 | 0.364 | 0.415 | 0.499 | 5000 |
| mca | 0.744 | 0.530 | 0.776 | 0.299 | 0.353 | 0.409 | 5000 |
| hcl | 0.850 | 0.581 | 0.505 | 0.368 | 0.426 | 0.488 | 5000 |
| immune | 1.207 | 0.802 | 0.644 | 0.552 | 0.608 | 0.655 | 5000 |
| cortex | 1.036 | 0.867 | 0.636 | 0.488 | 0.553 | 0.624 | 5000 |
| splatter1 | 1.145 | 1.135 | 0.990 | 0.911 | 0.950 | 0.985 | 10000 |
| splatter2 | 1.137 | 1.080 | 0.997 | 0.914 | 0.950 | 0.990 | 10000 |
| splatter3 | 1.117 | 1.156 | 0.997 | 0.916 | 0.950 | 0.989 | 10000 |
| splatter4 | 1.117 | 1.099 | 0.994 | 0.919 | 0.959 | 0.995 | 10000 |
| splatter5 | 1.139 | 1.077 | 0.989 | 0.914 | 0.953 | 0.993 | 10000 |
| splatter6 | 1.106 | 1.080 | 1.011 | 0.922 | 0.959 | 0.992 | 10000 |
| splatter7 | 1.109 | 1.051 | 1.020 | 0.937 | 0.973 | 1.008 | 10000 |
| splatter8 | 1.055 | 1.050 | 1.003 | 0.934 | 0.971 | 0.996 | 10000 |
| heart | 0.826 | 0.636 | 0.366 | 0.347 | 0.380 | 0.425 | 10000 |
| moffit | 1.099 | 0.783 | 0.566 | 0.539 | 0.581 | 0.636 | 10000 |
| fetal | 0.926 | 0.554 | 0.359 | 0.343 | 0.384 | 0.444 | 10000 |
| pbmc68k | 1.190 | 0.920 | 0.677 | 0.633 | 0.669 | 0.709 | 10000 |
| bacher | 1.454 | 0.906 | 0.765 | 0.696 | 0.738 | 0.785 | 10000 |
| zeisel | 1.655 | 0.751 | 0.360 | 0.322 | 0.359 | 0.412 | 10000 |
| mca | 0.725 | 0.511 | 0.324 | 0.259 | 0.298 | 0.354 | 10000 |
| hcl | 0.826 | 0.523 | 0.458 | 0.326 | 0.370 | 0.433 | 10000 |
| immune | 1.202 | 0.795 | 0.579 | 0.499 | 0.544 | 0.602 | 10000 |
| cortex | 1.036 | 0.717 | 0.547 | 0.442 | 0.497 | 0.564 | 10000 |

Table S4: Computation time

| dataset | random | Geosketch | Hopper | scSampler-sp1 | scSampler-sp4 | scSampler-sp16 | subsample size |
|---|---|---|---|---|---|---|---|
| splatter1 | 0.001 | 140.223 | 4.769 | 13.508 | 2.738 | 0.558 | 1000 |
| splatter2 | 0.000 | 140.002 | 4.771 | 13.422 | 2.686 | 0.560 | 1000 |
| splatter3 | 0.000 | 107.804 | 9.603 | 55.030 | 2.711 | 0.559 | 1000 |
| splatter4 | 0.000 | 105.439 | 6.454 | 14.177 | 2.768 | 0.564 | 1000 |
| splatter5 | 0.000 | 106.055 | 6.355 | 14.114 | 2.750 | 0.558 | 1000 |
| splatter6 | 0.000 | 213.694 | 8.905 | 30.130 | 5.831 | 1.069 | 1000 |
| splatter7 | 0.001 | 717.140 | 16.547 | 73.694 | 17.059 | 3.597 | 1000 |
| splatter8 | 0.001 | 1279.454 | 36.180 | 148.260 | 34.024 | 7.356 | 1000 |
| heart | 0.000 | 35.682 | 4.973 | 3.314 | 0.644 | 0.207 | 1000 |
| moffit | 0.000 | 36.392 | 5.072 | 3.449 | 0.664 | 0.211 | 1000 |
| fetal | 0.000 | 52.677 | 5.657 | 7.173 | 1.358 | 0.359 | 1000 |
| pbmc68k | 0.000 | 75.840 | 4.414 | 7.886 | 1.507 | 0.400 | 1000 |
| bacher | 0.000 | 128.754 | 5.815 | 12.052 | 2.934 | 0.582 | 1000 |
| zeisel | 0.000 | 172.442 | 5.578 | 18.269 | 4.743 | 0.880 | 1000 |
| mca | 0.000 | 437.237 | 11.498 | 51.094 | 9.778 | 2.154 | 1000 |
| hcl | 0.001 | 762.103 | 16.867 | 88.803 | 16.890 | 4.369 | 1000 |
| immune | 0.001 | 441.763 | 21.736 | 90.124 | 21.027 | 4.480 | 1000 |
| cortex | 0.001 | 1396.994 | 32.271 | 164.974 | 41.933 | 7.982 | 1000 |
| splatter1 | 0.000 | 106.108 | 12.182 | 34.120 | 8.174 | 1.651 | 3000 |
| splatter2 | 0.000 | 107.679 | 14.099 | 34.194 | 8.186 | 1.654 | 3000 |
| splatter3 | 0.000 | 145.947 | 14.976 | 34.573 | 8.250 | 1.659 | 3000 |
| splatter4 | 0.000 | 141.908 | 15.302 | 34.518 | 8.188 | 1.656 | 3000 |
| splatter5 | 0.000 | 142.565 | 13.328 | 34.234 | 8.234 | 1.643 | 3000 |
| splatter6 | 0.000 | 306.005 | 14.908 | 90.995 | 17.356 | 3.146 | 3000 |
| splatter7 | 0.000 | 632.163 | 30.011 | 223.600 | 42.275 | 10.824 | 3000 |
| splatter8 | 0.001 | 1459.032 | 37.614 | 491.348 | 107.120 | 20.997 | 3000 |
| heart | 0.000 | 31.116 | 14.103 | 9.284 | 1.860 | 0.600 | 3000 |
| moffit | 0.000 | 26.246 | 14.847 | 9.667 | 1.903 | 0.611 | 3000 |
| fetal | 0.000 | 64.507 | 10.538 | 20.693 | 4.017 | 1.064 | 3000 |
| pbmc68k | 0.000 | 85.686 | 9.069 | 22.852 | 4.711 | 1.174 | 3000 |
| bacher | 0.000 | 111.752 | 14.536 | 34.902 | 8.545 | 1.723 | 3000 |
| zeisel | 0.000 | 62.611 | 11.908 | 65.994 | 14.100 | 2.576 | 3000 |
| mca | 0.001 | 423.334 | 14.709 | 155.319 | 39.714 | 8.496 | 3000 |
| hcl | 0.001 | 217.882 | 22.179 | 259.642 | 59.929 | 12.675 | 3000 |
| immune | 0.001 | 762.271 | 26.606 | 269.259 | 61.348 | 12.737 | 3000 |
| cortex | 0.001 | 1190.904 | 37.704 | 519.825 | 117.953 | 22.737 | 3000 |
| splatter1 | 0.000 | 140.947 | 22.572 | 65.946 | 13.641 | 2.731 | 5000 |
| splatter2 | 0.000 | 144.306 | 22.948 | 54.999 | 13.245 | 2.750 | 5000 |
| splatter3 | 0.000 | 125.375 | 21.389 | 54.737 | 13.222 | 2.753 | 5000 |
| splatter4 | 0.000 | 124.000 | 16.760 | 54.836 | 13.211 | 2.738 | 5000 |
| splatter5 | 0.000 | 144.177 | 16.994 | 54.526 | 13.191 | 2.730 | 5000 |
| splatter6 | 0.000 | 287.245 | 19.211 | 148.592 | 28.135 | 5.295 | 5000 |
| splatter7 | 0.000 | 541.812 | 28.854 | 374.113 | 85.754 | 17.549 | 5000 |
| splatter8 | 0.001 | 1457.772 | 47.872 | 725.685 | 182.585 | 35.120 | 5000 |
| heart | 0.000 | 25.621 | 17.085 | 15.094 | 3.016 | 0.971 | 5000 |
| moffit | 0.000 | 32.226 | 16.199 | 15.741 | 3.122 | 0.993 | 5000 |
| fetal | 0.000 | 53.684 | 16.726 | 33.273 | 6.554 | 1.748 | 5000 |
| pbmc68k | 0.000 | 84.608 | 11.755 | 37.046 | 9.294 | 2.930 | 5000 |
| bacher | 0.000 | 111.618 | 16.787 | 66.491 | 13.999 | 2.837 | 5000 |

| | | | | | | | |
|---|---|---|---|---|---|---|---|
| zeisel | 0.000 | 174.142 | 14.675 | 106.698 | 22.809 | 4.255 | 5000 |
| mca | 0.000 | 421.410 | 16.965 | 242.229 | 45.774 | 10.230 | 5000 |
| hcl | 0.001 | 648.598 | 23.462 | 453.807 | 82.780 | 21.087 | 5000 |
| immune | 0.001 | 863.776 | 26.138 | 439.458 | 100.104 | 21.005 | 5000 |
| cortex | 0.001 | 1577.870 | 41.540 | 789.105 | 203.926 | 38.887 | 5000 |
| splatter1 | 0.000 | 144.649 | 53.135 | 142.018 | 25.545 | 5.388 | 10000 |
| splatter2 | 0.000 | 125.913 | 31.183 | 133.151 | 25.546 | 5.379 | 10000 |
| splatter3 | 0.001 | 125.804 | 33.334 | 156.299 | 25.060 | 5.332 | 10000 |
| splatter4 | 0.000 | 125.849 | 30.728 | 132.935 | 25.014 | 5.358 | 10000 |
| splatter5 | 0.000 | 126.760 | 30.822 | 131.738 | 28.447 | 7.021 | 10000 |
| splatter6 | 0.000 | 283.663 | 43.516 | 286.487 | 55.147 | 10.571 | 10000 |
| splatter7 | 0.001 | 737.263 | 101.804 | 725.448 | 169.660 | 34.494 | 10000 |
| splatter8 | 0.001 | 1267.002 | 105.263 | 1503.680 | 362.096 | 69.903 | 10000 |
| heart | 0.000 | 42.044 | 42.337 | 30.133 | 5.748 | 1.879 | 10000 |
| moffit | 0.000 | 38.251 | 43.735 | 29.993 | 5.911 | 1.936 | 10000 |
| fetal | 0.000 | 94.284 | 68.868 | 65.173 | 12.482 | 3.379 | 10000 |
| pbmc68k | 0.000 | 84.653 | 21.709 | 72.842 | 14.855 | 3.758 | 10000 |
| bacher | 0.000 | 148.804 | 31.699 | 139.489 | 27.066 | 5.545 | 10000 |
| zeisel | 0.000 | 234.393 | 27.070 | 208.887 | 44.469 | 8.399 | 10000 |
| mca | 0.001 | 493.098 | 33.987 | 477.034 | 109.250 | 33.801 | 10000 |
| hcl | 0.001 | 539.579 | 38.243 | 854.913 | 217.309 | 42.359 | 10000 |
| immune | 0.001 | 768.613 | 37.541 | 881.611 | 201.097 | 42.817 | 10000 |
| cortex | 0.001 | 1389.987 | 49.148 | 1652.981 | 456.728 | 78.101 | 10000 |

Table S5: Gini coefficients in subsamples

| dataset | random | Geosketch | Hopper | scSampler-sp1 | scSampler-sp4 | scSampler-sp16 | subsample size |
|---|---|---|---|---|---|---|---|
| splatter1 | 0.495 | 0.398 | 0.384 | 0.398 | 0.401 | 0.343 | 1000 |
| splatter2 | 0.512 | 0.407 | 0.419 | 0.359 | 0.352 | 0.313 | 1000 |
| splatter3 | 0.512 | 0.418 | 0.378 | 0.395 | 0.382 | 0.355 | 1000 |
| splatter4 | 0.512 | 0.383 | 0.380 | 0.356 | 0.352 | 0.294 | 1000 |
| splatter5 | 0.512 | 0.380 | 0.375 | 0.388 | 0.367 | 0.311 | 1000 |
| splatter6 | 0.484 | 0.421 | 0.418 | 0.369 | 0.356 | 0.344 | 1000 |
| splatter7 | 0.518 | 0.360 | 0.349 | 0.332 | 0.350 | 0.293 | 1000 |
| splatter8 | 0.502 | 0.371 | 0.391 | 0.368 | 0.361 | 0.310 | 1000 |
| heart | 0.525 | 0.500 | 0.356 | 0.393 | 0.383 | 0.397 | 1000 |
| moffit | 0.766 | 0.410 | 0.507 | 0.457 | 0.440 | 0.423 | 1000 |
| fetal | 0.644 | 0.415 | 0.396 | 0.426 | 0.383 | 0.324 | 1000 |
| pbmc68k | 0.526 | 0.647 | 0.629 | 0.702 | 0.712 | 0.688 | 1000 |
| bacher | 0.582 | 0.350 | 0.340 | 0.391 | 0.378 | 0.404 | 1000 |
| zeisel | 0.665 | 0.680 | 0.461 | 0.525 | 0.487 | 0.479 | 1000 |
| mca | 0.587 | 0.665 | 0.602 | 0.640 | 0.601 | 0.538 | 1000 |
| hcl | 0.497 | 0.753 | 0.620 | 0.632 | 0.606 | 0.599 | 1000 |
| immune | 0.569 | 0.759 | 0.679 | 0.731 | 0.708 | 0.672 | 1000 |
| cortex | 0.747 | 0.727 | 0.502 | 0.567 | 0.550 | 0.549 | 1000 |
| splatter1 | 0.510 | 0.437 | 0.405 | 0.420 | 0.405 | 0.393 | 3000 |
| splatter2 | 0.489 | 0.437 | 0.408 | 0.387 | 0.381 | 0.361 | 3000 |
| splatter3 | 0.489 | 0.456 | 0.382 | 0.416 | 0.402 | 0.385 | 3000 |
| splatter4 | 0.489 | 0.410 | 0.404 | 0.369 | 0.368 | 0.344 | 3000 |
| splatter5 | 0.489 | 0.426 | 0.400 | 0.392 | 0.380 | 0.358 | 3000 |
| splatter6 | 0.480 | 0.436 | 0.387 | 0.383 | 0.377 | 0.378 | 3000 |
| splatter7 | 0.506 | 0.425 | 0.370 | 0.370 | 0.366 | 0.334 | 3000 |
| splatter8 | 0.501 | 0.444 | 0.385 | 0.381 | 0.388 | 0.350 | 3000 |
| heart | 0.527 | 0.420 | 0.405 | 0.416 | 0.399 | 0.374 | 3000 |
| moffit | 0.770 | 0.519 | 0.542 | 0.522 | 0.514 | 0.498 | 3000 |
| fetal | 0.640 | 0.399 | 0.458 | 0.488 | 0.444 | 0.393 | 3000 |
| pbmc68k | 0.551 | 0.580 | 0.631 | 0.663 | 0.668 | 0.659 | 3000 |
| bacher | 0.587 | 0.366 | 0.357 | 0.392 | 0.387 | 0.381 | 3000 |
| zeisel | 0.648 | 0.596 | 0.473 | 0.542 | 0.497 | 0.464 | 3000 |
| mca | 0.566 | 0.664 | 0.584 | 0.654 | 0.618 | 0.579 | 3000 |
| hcl | 0.493 | 0.705 | 0.585 | 0.650 | 0.617 | 0.600 | 3000 |
| immune | 0.570 | 0.710 | 0.684 | 0.745 | 0.723 | 0.690 | 3000 |
| cortex | 0.736 | 0.679 | 0.592 | 0.595 | 0.567 | 0.543 | 3000 |
| splatter1 | 0.506 | 0.440 | 0.414 | 0.427 | 0.412 | 0.408 | 5000 |
| splatter2 | 0.491 | 0.433 | 0.426 | 0.402 | 0.396 | 0.379 | 5000 |
| splatter3 | 0.491 | 0.451 | 0.404 | 0.417 | 0.415 | 0.400 | 5000 |
| splatter4 | 0.491 | 0.420 | 0.400 | 0.383 | 0.383 | 0.364 | 5000 |
| splatter5 | 0.491 | 0.404 | 0.431 | 0.400 | 0.391 | 0.382 | 5000 |
| splatter6 | 0.480 | 0.438 | 0.378 | 0.398 | 0.388 | 0.385 | 5000 |
| splatter7 | 0.503 | 0.404 | 0.395 | 0.382 | 0.379 | 0.361 | 5000 |
| splatter8 | 0.497 | 0.428 | 0.389 | 0.381 | 0.379 | 0.358 | 5000 |
| heart | 0.530 | 0.399 | 0.402 | 0.423 | 0.413 | 0.389 | 5000 |
| moffit | 0.766 | 0.583 | 0.599 | 0.565 | 0.559 | 0.553 | 5000 |
| fetal | 0.637 | 0.399 | 0.491 | 0.514 | 0.471 | 0.429 | 5000 |
| pbmc68k | 0.551 | 0.545 | 0.591 | 0.631 | 0.633 | 0.636 | 5000 |
| bacher | 0.585 | 0.364 | 0.382 | 0.399 | 0.391 | 0.390 | 5000 |

| | | | | | | | |
|---|---|---|---|---|---|---|---|
| zeisel | 0.646 | 0.551 | 0.505 | 0.546 | 0.509 | 0.473 | 5000 |
| mca | 0.569 | 0.667 | 0.594 | 0.662 | 0.631 | 0.599 | 5000 |
| hcl | 0.491 | 0.689 | 0.589 | 0.652 | 0.624 | 0.602 | 5000 |
| immune | 0.565 | 0.706 | 0.697 | 0.756 | 0.735 | 0.705 | 5000 |
| cortex | 0.733 | 0.658 | 0.595 | 0.607 | 0.580 | 0.555 | 5000 |
| splatter1 | 0.496 | 0.448 | 0.427 | 0.433 | 0.430 | 0.423 | 10000 |
| splatter2 | 0.498 | 0.439 | 0.434 | 0.417 | 0.412 | 0.406 | 10000 |
| splatter3 | 0.498 | 0.445 | 0.437 | 0.430 | 0.424 | 0.420 | 10000 |
| splatter4 | 0.498 | 0.410 | 0.422 | 0.404 | 0.399 | 0.388 | 10000 |
| splatter5 | 0.498 | 0.405 | 0.436 | 0.414 | 0.407 | 0.397 | 10000 |
| splatter6 | 0.489 | 0.430 | 0.401 | 0.407 | 0.401 | 0.399 | 10000 |
| splatter7 | 0.494 | 0.400 | 0.398 | 0.397 | 0.394 | 0.382 | 10000 |
| splatter8 | 0.492 | 0.418 | 0.404 | 0.389 | 0.387 | 0.371 | 10000 |
| heart | 0.538 | 0.424 | 0.437 | 0.441 | 0.430 | 0.414 | 10000 |
| moffit | 0.768 | 0.655 | 0.656 | 0.635 | 0.633 | 0.634 | 10000 |
| fetal | 0.628 | 0.423 | 0.513 | 0.536 | 0.507 | 0.471 | 10000 |
| pbmc68k | 0.544 | 0.484 | 0.527 | 0.563 | 0.562 | 0.559 | 10000 |
| bacher | 0.575 | 0.359 | 0.396 | 0.400 | 0.392 | 0.389 | 10000 |
| zeisel | 0.646 | 0.512 | 0.540 | 0.561 | 0.530 | 0.491 | 10000 |
| mca | 0.567 | 0.658 | 0.637 | 0.675 | 0.645 | 0.609 | 10000 |
| hcl | 0.482 | 0.667 | 0.596 | 0.656 | 0.632 | 0.609 | 10000 |
| immune | 0.574 | 0.706 | 0.703 | 0.762 | 0.743 | 0.717 | 10000 |
| cortex | 0.732 | 0.640 | 0.609 | 0.618 | 0.600 | 0.573 | 10000 |

### References


[Bacher et al., 2020] Bacher, P., Rosati, E., Esser, D., Martini, G. R., Saggau, C., Schiminsky, E., Dargvainiene, J., Schröder, I., Wieters, I., Khodamoradi, Y., et al. (2020). Low-avidity cd4+ t cell responses to sars-cov-2 in unexposed individuals and humans with severe covid-19. *Immunity*, 53(6):1258–1271.

[Han et al., 2018] Han, X., Wang, R., Zhou, Y., Fei, L., Sun, H., Lai, S., Saadatpour, A., Zhou, Z., Chen, H., Ye, F., et al. (2018). Mapping the mouse cell atlas by microwell-seq. *Cell*, 172(5):1091–1107.

[Han et al., 2020] Han, X., Zhou, Z., Fei, L., Sun, H., Wang, R., Chen, Y., Chen, H., Wang, J., Tang, H., Ge, W., et al. (2020). Construction of a human cell landscape at single-cell level. *Nature*, 581(7808):303–309.

[McLellan et al., 2020] McLellan, M. A., Skelly, D. A., Dona, M. S., Squiers, G. T., Farrugia, G. E., Gaynor, T. L., Cohen, C. D., Pandey, R., Diep, H., Vinh, A., et al. (2020). High-resolution transcriptomic profiling of the heart during chronic stress reveals cellular drivers of cardiac fibrosis and hypertrophy. *Circulation*, 142(15):1448–1463.

[Moffitt et al., 2018] Moffitt, J. R., Bambah-Mukku, D., Eichhorn, S. W., Vaughn, E., Shekhar, K., Perez, J. D., Rubinstein, N. D., Hao, J., Regev, A., Dulac, C., et al. (2018). Molecular, spatial, and functional single-cell profiling of the hypothalamic preoptic region. *Science*, 362(6416).

[Papatheodorou et al., 2020] Papatheodorou, I., Moreno, P., Manning, J., Fuentes, A. M.-P., George, N., Fexova, S., Fonseca, N. A., Füllgrabe, A., Green, M., Huang, N., et al. (2020). Expression atlas update: from tissues to single cells. *Nucleic acids research*, 48(D1):D77–D83.

[Vento-Tormo et al., 2018] Vento-Tormo, R., Efremova, M., Botting, R. A., Turco, M. Y., Vento-Tormo, M., Meyer, K. B., Park, J.-E., Stephenson, E., Polański, K., Goncalves, A., et al. (2018). Single-cell reconstruction of the early maternal–fetal interface in humans. *Nature*, 563(7731):347–353.

[Yao et al., 2021] Yao, Z., van Velthoven, C. T., Nguyen, T. N., Goldy, J., Sedeno-Cortes, A. E., Baftizadeh, F., Bertagnolli, D., Casper, T., Chiang, M., Crichton, K., et al. (2021). A taxonomy of transcriptomic cell types across the isocortex and hippocampal formation. *Cell*, 184(12):3222–3241.

[Zappia et al., 2017] Zappia, L., Phipson, B., and Oshlack, A. (2017). Splatter: simulation of single-cell rna sequencing data. *Genome biology*, 18(1):1–15.

[Zeisel et al., 2015] Zeisel, A., Muñoz-Manchado, A. B., Codeluppi, S., Lönnerberg, P., La Manno, G., Juréus, A., Marques, S., Munguba, H., He, L., Betsholtz, C., et al. (2015). Cell types in the mouse cortex and hippocampus revealed by single-cell rna-seq. *Science*, 347(6226):1138–1142.

[Zheng et al., 2017] Zheng, G. X., Terry, J. M., Belgrader, P., Ryvkin, P., Bent, Z. W., Wilson, R., Ziraldo, S. B., Wheeler, T. D., McDermott, G. P., Zhu, J., et al. (2017). Massively parallel digital transcriptional profiling of single cells. *Nature communications*, 8(1):1–12.
